# A kinesin mediates VEGFR2 recycling and regulates VE-cadherin phosphorylation, essential for vascular permeability

**DOI:** 10.1101/2022.06.06.494986

**Authors:** Hyun-Dong Cho, Christopher Zhou, Kayeman Tu, Tara Nguyen, Nicolene A. Sarich, Kaori H. Yamada

**Author notes:** Corresponding author. Kaori H. Yamada.

## Abstract

Excessive vascular endothelial growth factor-A (VEGF-A) signaling induces vascular leakage and angiogenesis in diseases. VEGFR2 trafficking to the cell surface, mediated by kinesin-3 family protein KIF13B, is essential to respond to VEGF-A in inducing angiogenesis. However, the precise mechanism of how KIF13B regulates VEGF-induced signaling and endothelial permeability is unknown. Here we show that KIF13B-mediated recycling of internalized VEGFR2 through Rab11-positive recycling vesicle regulates VE-cadherin phosphorylation and endothelial permeability. Phosphorylated VEGFR2 at the cell-cell junction was internalized and associated with KIF13B in Rab5-positive early endosomes. KIF13B mediated VEGFR2 recycling through Rab11-positive recycling vesicle, and inhibition of this recycling attenuated phosphorylation of VEGFR2 at Y951, Src, and VE-cadherin at Y685, which are necessary for endothelial permeability. Failure of VEGFR2 trafficking to the cell surface induced accumulation and degradation of VEGFR2 in lysosomes. Furthermore, in the animal model of wet age-related macular degeneration (AMD), inhibition of KIF13B-mediated VEGFR2 trafficking also mitigated vascular leakage. Thus, the present results identify the fundamental role of VEGFR2 recycling to the cell surface in mediating vascular permeability, suggesting a promising strategy for mitigating vascular leakage associated with inflammatory diseases.

## Introduction

Leakage from blood vessels at the level of endothelial cell junctions causes tissue inflammation in many diseases, including cancer and blinding eye diseases (Claesson-Welsh et al., 2020). Endothelial cells lining the vessel wall have adherens junctions and tight junctions, which serve as barriers limiting the permeation of molecules and liquid (Claesson-Welsh et al., 2020; Duong and Vestweber, 2020). In diseased conditions, excessive vascular endothelial growth factor-A (VEGF-A) induces unchecked angiogenesis and vascular leakage (Claesson-Welsh et al., 2020; Eelen et al., 2020; Apte et al., 2019). We have developed a new strategy to inhibit pathological angiogenesis in cancer (Yamada et al., 2017) and wet age-related macular degeneration (wet AMD) (Waters et al., 2021b), a leading cause of blindness. However, whether this approach has an anti-endothelial permeability function remains unknown. Here we show that inhibition of VEGFR2 trafficking to the plasma membrane was critical in controlling endothelial permeability.

The cell surface receptor for VEGF-A, VEGFR2, is trafficked between the cell surface and intra-endothelial domains (Simons et al., 2016). VEGFR2 in the quiescent endothelium is normally present in a dephosphorylated inactive state on the endothelial cell surface (Simons et al., 2016). Upon binding to VEGF-A, phosphorylation of VEGFR2 rapidly induces internalization (Nakayama et al., 2013; Sawamiphak et al., 2010) and trafficking into early endosomes and its colocalization with Rab5 (Lanahan et al., 2010). VEGFR2 is subsequently dephosphorylated (Lanahan et al., 2014) before entering Rab11 recycling vesicles (Ballmer-Hofer et al., 2011). Part of the VEGFR2 pool is sorted for recycling to the cell surface (Simons et al., 2016) to make the receptor available for another round of ligand binding (Lanahan et al., 2010; Manickam et al., 2011; Tiwari et al., 2013). The non-cycled part of VEGFR2 undergoes degradation through Rab7-positive late endosomes and lysosomal membrane-associated protein 2 (LAMP2)-positive lysosome (Simons et al., 2016). To restore VEGFR2 levels on the plasmalemma, newly synthesized VEGFR2 is trafficked from the Golgi apparatus to the cell surface (Manickam et al., 2011; Yamada et al., 2014). We found that trafficking of VEGFR2 from the Golgi apparatus to the cell surface is mediated by a kinesin-3 family molecular motor KIF13B (Yamada et al., 2014). KIF13B knockout mice (*Kif13b*^*KO*^) showed reduced angiogenesis (Waters et al., 2021b). Endothelial-specific inducible KIF13B knockout (*Kif13b*^*iECKO*^) also showed reduced angiogenesis, vascular leakage, and cancer metastasis (Waters et al., 2021a). Furthermore, pharmacological disruption of the interaction between KIF13B and VEGFR2 inhibited vascular leakage (Waters et al., 2021a) and neovascularization (Waters et al., 2021b).

Although VEGF-A induces angiogenesis and endothelial permeability, the signaling pathways are different (Simons et al., 2016; Claesson-Welsh et al., 2020). Phosphorylation of VEGFR2 at Y951 recruits T cell-specific adaptor (TSAd), which regulates activation of c-Src tyrosine kinase (Smith et al., 2020). Src-mediated phosphorylation of Y685 of VE-cadherin is essential to increase endothelial permeability by VEGF-A (Orsenigo et al., 2012; Adam et al., 2010; Wessel et al., 2014). Although the importance of VEGFR2 trafficking is well known, we surmised whether VEGFR2 trafficking might also regulate endothelial permeability, a relatively rapid response occurring within minutes (Claesson-Welsh et al., 2020) as contrasted with angiogenesis requiring hours (Simons et al., 2016). Here we addressed mechanisms regulating VEGF-A induced endothelial permeability. Among the approaches employed was the peptide KAI (kinesin-derived angiogenesis inhibitor), which inhibits KIF13B-mediated VEGFR2 trafficking (Yamada et al., 2017) as well as knockdown of the molecular motor KIF13B in endothelial cells that traffics the receptor to the plasma membrane. We showed the crucial role of KIF13B mediated VEGFR2 trafficking to plasma membrane in regulating VEGF-A endothelial permeability.

## Results

### KIF13B regulates VEGF-induced endothelial permeability

KIF13B transports newly synthesized VEGFR2 from Golgi to the cell surface (Yamada et al., 2014), and the VEGFR2 trafficking is essential for angiogenesis (Waters et al., 2021b; Yamada et al., 2017, 2014). We showed that KIF13B also plays an important role in VEGF-induced vascular leakage, as VEGF-A fails to induce vascular leakage in *Kif13b*^*KO*^ or when KIF13B-mediated VEGFR2 trafficking is blocked by KAI (Waters et al., 2021a). However, VEGF-induced endothelial permeability is an early response of ECs to VEGF-A, occurring within 15 min. It cannot be explained by the known function of KIF13B, which is the release of synthesized VEGFR2 from Golgi and its trafficking to the cell surface, taking several hours after VEGF-A stimulation (Yamada et al., 2014). To explore whether KIF13B has any role in the relatively rapid response to VEGF-A, we used lentivirus-based shRNA-*KIF13B* to knockdown (Yamada et al., 2014), and tested VEGF-induced endothelial permeability in human retina endothelial cells (hREC) using transendothelial electric resistance (TEER) assay. VEGF-A transiently reduced TEER at 15 min after VEGF-A stimulation in control hREC, transduced with scrambled shRNA (**Fig. 1A, B**), whereas knockdown of *KIF13B* suppressed VEGF-induced endothelial permeability compared to controls (**Fig. 1A, B**). After initial permeability, the endothelial barrier function recovered in both scrambled and shRNA-*KIF13B* treated cells (**Fig. 1A, C**).

**Fig. 1.**
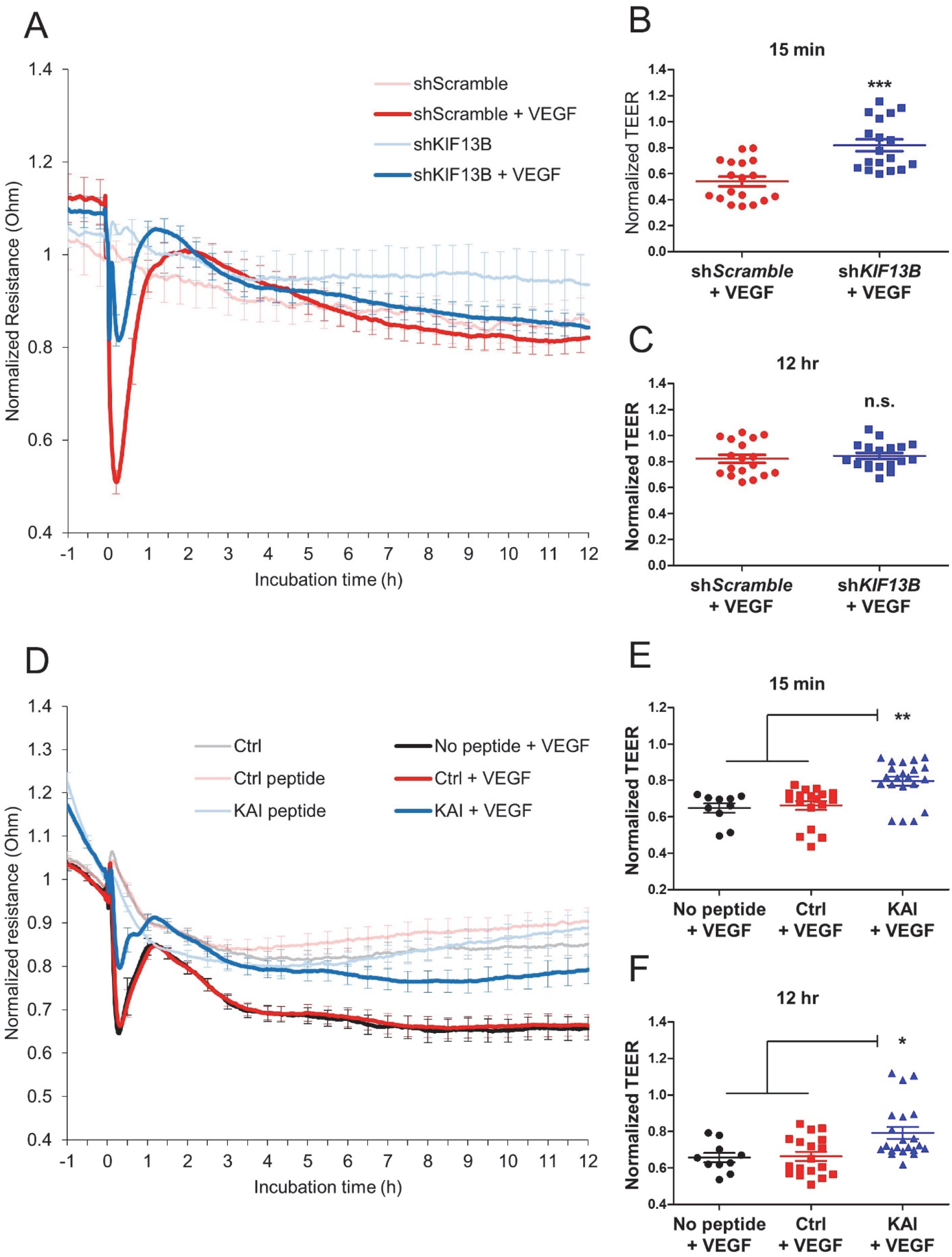
KIF13B is essential for VEGF-induced permeability in hREC. **(A, B, C)** Transendothelial electrical resistance (TEER) measurement of confluent hRECs transduced with scrambled shRNA (red and pink lines) or shRNA-*KIF13B* (blue and light blue) expressed as normalized resistance of the TEER basal values. Cells were stimulated with VEGF-A (50 ng/mL) at time zero, and TEER was measured every 30 seconds over time (4,000 Hz of frequency). Normalized resistance changes overtime were shown in graph A. Normalized resistance at 15 min after VEGF-A stimulation and 12 hr after VEGF-A stimulation were shown in B, and C as mean ± SE. N=5, 6, 18, 18, for scrambled shRNA, shRNA-*KIF13B*, scrambled shRNA + VEGF-A, and shRNA-*KIF13B* + VEGF-A, respectively. One-way ANOVA with post hoc multiple comparisons. *** indicates p<0.001. **(D, E, F)** TEER measurements of hREC treated with KAI (10 µM), an inhibitor for VEGFR2 trafficking, or control peptide (10 µM) followed by VEGF-A (50 ng/mL) stimulation. Changes of normalized resistance over time were shown in graph D. Normalized resistance at 15 min and 12 hr after VEGF-A stimulation were shown in E and F as mean ± SE. N=12, 10, 13, 8, 16, 15, for no peptide, ctrl, KAI, no peptide + VEGF-A, ctrl + VEGF-A, KAI + VEGF-A, respectively. One-way ANOVA with post hoc multiple comparisons. * indicates p<0.05, ** indicates p<0.01.

Next, we tested whether the inhibition of VEGFR2 trafficking also affects endothelial permeability (**Fig. 1D, E, F**). KAI is a synthetic peptide designed to disrupt the interaction between VEGFR2 and KIF13B (Yamada et al., 2017). After pretreatment with the control peptide or KAI (10 µM) for 2 hours, the TEER of confluent hREC was monitored before and after VEGF-A stimulation (**Fig. 1D**). KAI treatment prevented VEGF-induced endothelial permeability, compared to no-peptide control and control peptide treated hREC (**Fig. 1D**). KAI-treated hREC showed higher electrical resistance values than both no peptide and control peptide-treated cells 15 min and 12 hr after stimulation with VEGF-A (**Fig. 1D, E, F**). These data demonstrate that KIF13B is required for VEGF-induced endothelial permeability.

### KIF13B regulates VEGFR2 signaling pathways mediating endothelial permeability

To examine how KIF13B regulates endothelial permeability, we examined the activation of the signaling pathway regulating endothelial permeability and tested the effect of knockdown of *KIF13B* (**Fig. 2A, B**). VEGF-A treatment increased phosphorylation of VEGFR2 (at Y951 and Y1175), Src (at Y416), and VE-cadherin (at Y658 and Y685) in scrambled shRNA transduced control hREC. Compared to scrambled shRNA transduced cells, shRNA-*KIF13B* transduced cells showed significantly lower phosphorylated/total protein ratio in p-VEGFR2 (Y951 and Y1175), p-Src (Y416), and p-VE-cadherin (Y658 and Y685) through 3-15 min of VEGF-A treatment (**Fig. 2A, B**).

**Fig. 2.**
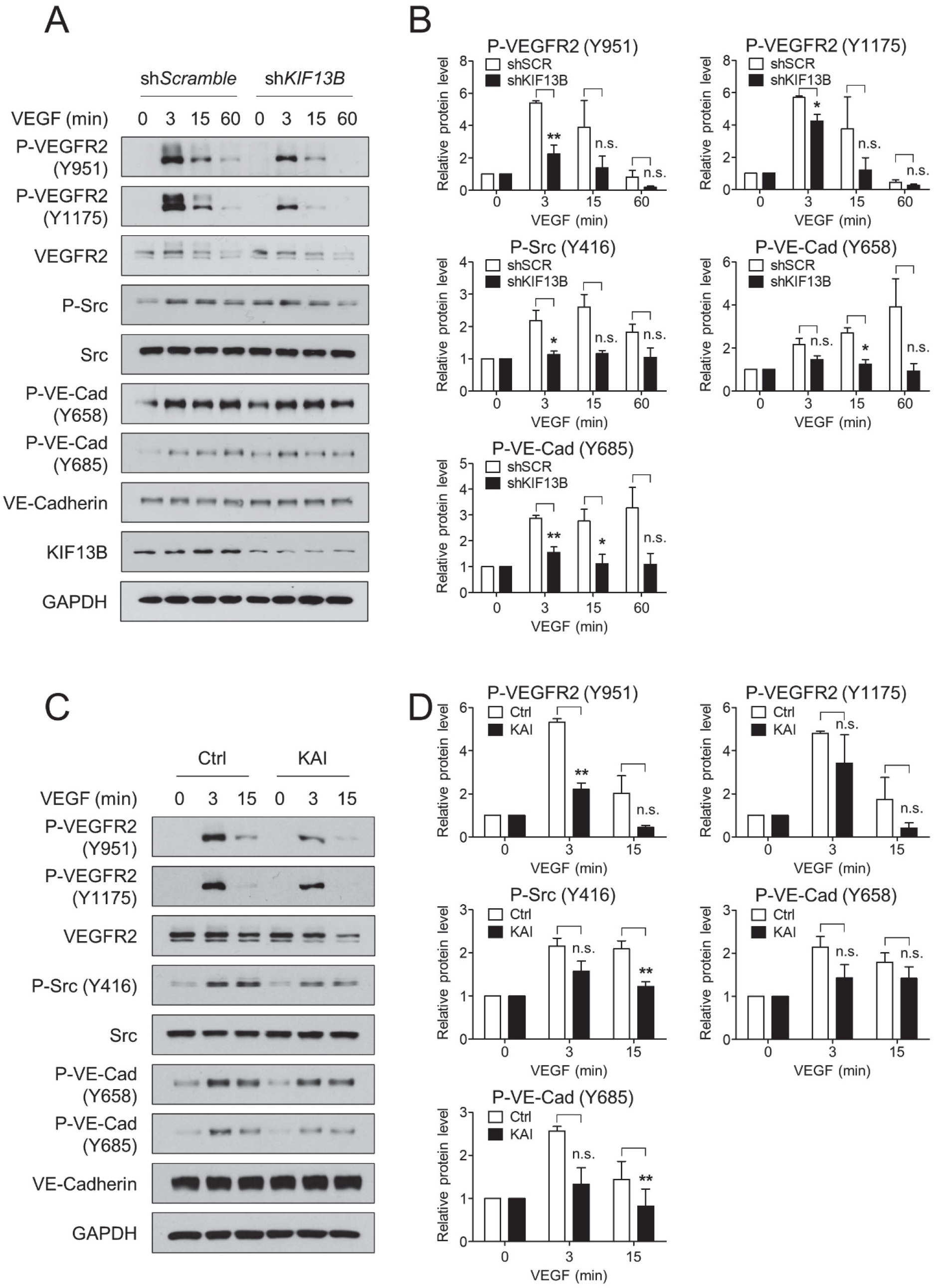
KIF13B is required to regulate VEGFR2 signaling pathway. **(A, B)** VEGF-induced phosphorylation of signaling molecules in hREC transduced with either scrambled shRNA or shRNA-*KIF13B*. After stimulation with VEGF-A (50 ng/mL) for indicated time points, cell lysates were analyzed by western blotting for p-VEGFR2 (Y951), p-VEGFR2 (Y1175), VEGFR2, p-Src (S416), Src, p-VE-cadherin (Y658), p-VE-cadherin (Y685), VE-cadherin, KIF13B, and GAPDH (p indicates the phosphorylated form). Representative blots were shown in A. Quantification of blots of phosphorylated proteins expressed relative to total proteins were shown as mean ± SE in graph B. N=3. One-way ANOVA followed by post hoc multiple comparisons. * indicates p<0.05, ** indicates p<0.01. No significant difference (n.s.) indicates p>0.05. **(C, D)** VEGF-induced phosphorylation of signaling molecules in hREC treated with either control peptide or KAI (10 µM). After stimulation with VEGF-A (50 ng/mL) for indicated time points, cell lysates were analyzed by western blotting for phosphorylated and total proteins. Representative blots were shown in C. Quantification of phosphorylated proteins relative to total proteins was shown as mean ± SE in graph B. N=4. One-way ANOVA followed by post hoc multiple comparisons. ** indicates p<0.01. n.s. indicates p>0.05.

Next, we tested the effect of KAI compared with the control peptide (**Fig. 2C, D**). Similar to scrambled shRNA treated control above, VEGF-A induced p-VEGFR2 (at Y951 and Y1175), p-Src (at Y416), and p-VE-cadherin (at Y658 and Y685) in control peptide treated hREC (**Fig. 2C, D**). KAI treatment significantly decreased phosphorylation of VEGFR2 (Y951), Src (Y416), and VE-cadherin (Y685) compared with control peptide (**Fig. 2C, D**). KAI treatment also decreased p-VEGFR2 (Y1175) and pY658-VE-cadherin, although the difference was not statistically significant. Together, these data suggest the important role of KIF13B in regulating VEGF/VEGFR2 signaling to regulate endothelial permeability.

### KIF13B regulates translocation of internalized VEGFR2

As KIF13B mediates VEGFR2 trafficking to the cell surface (Yamada et al., 2014), we hypothesized that KIF13B tunes the degree of VEGF/VEGFR2 signaling by regulating the amount of VEGFR2 on the cell surface. We examined the effect of the knockdown of *KIF13B* on cell surface VEGFR2 before and after VEGF-A stimulation (**Fig. 3**). First, hREC were transduced with scrambled shRNA or shRNA-*KIF13B*, and stimulated with VEGF-A for indicated times. Cell surface proteins were biotinylated and precipitated with streptavidin beads, and biotinylated cell surface VEGFR2 was detected by western blotting (**Fig. 3A, B**). Cell surface VEGFR2 was decreased at 15-30 min after VEGF-A stimulation in both control and shRNA-*KIF13B* transduced hRECs, indicating VEGF-induced internalization (**Fig. 3A, B**). Interestingly, the knockdown of *KIF13B* further decreased the amount of VEGFR2 on the cell surface (**Fig. 3B**).

**Fig. 3.**
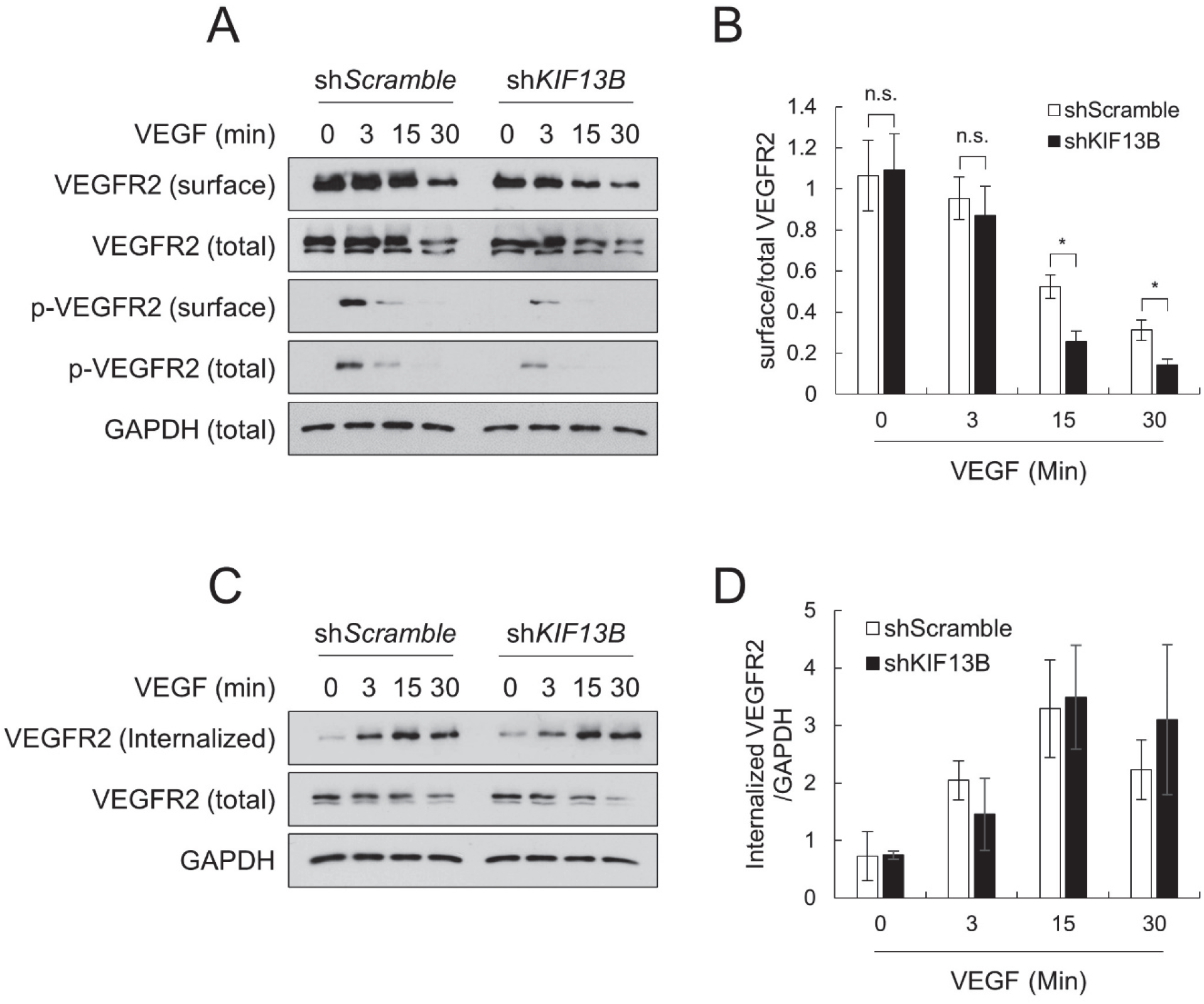
KIF13B deficiency reduces cell-surface VEGFR2 after VEGF stimulation. **(A, B)** Cell surface VEGFR2 levels detected by cell surface biotinylation of hREC transduced with either scrambled shRNA or shRNA-*KIF13B*, followed by VEGF-A stimulation (50 ng/mL) for indicated time periods. Total lysates (total) and streptavidin pull-down (surface), western blotting for VEGFR, p-VEGFR2 (Y1175) were shown in A. GAPDH was used as a loading control. Quantification of cell surface VEGFR2 was normalized by total VEGFR2, and shown as mean ± SE in graph B. N=4. One-way ANOVA followed by multiple comparisons. * indicates p<0.05. n.s. indicates p>0.05. **(C, D)** The internalized pool of VEGFR2 after VEGF-A treatment (50 ng/mL) of hREC transduced with either scrambled shRNA or shRNA-*KIF13B*. Cell surface biotinylation was performed prior to VEGF-A stimulation. At indicated time points, the remaining cell surface biotin was removed, and the internalized pool of VEGFR2 was collected by streptavidin pull-down. Western blotting of the total lysate (total) and streptavidin pull-down (internalized) for VEGFR2 was shown in C. Internalized VEGFR2 was normalized by loading control GAPDH, and shown as mean ± SE in the graph D. N=3. One-way ANOVA followed by multiple comparisons. * indicates p<0.05. n.s. indicates p>0.05.

As a component of internalized VEGFR2 is recycled back to the cell surface (Simons et al., 2016), the reduction of cell-surface VEGFR2 may be the result of increased internalization or less recycling. Thus, we used another biotinylation assay (Nitzsche et al., 2021) to quantify internalized VEGFR2 (**Fig. 3C, D**). hRECs were transduced with scrambled shRNA or shRNA-*KIF13B*. After the serum-starvation of cells, cell surface proteins were biotinylated. Then, cells were stimulated with VEGF-A to let VEGFR2 internalize. After removal of cell surface biotin, internalized VEGFR2 was detected by western blotting in streptavidin precipitates (**Fig. 3C, D**). After VEGF-A stimulation, VEGFR2 was internalized in both control and shRNA-*KIF13B* transduced cells without any difference (**Fig. 3C, D**), suggesting the possibility of KIF13B mediating recycling of internalized VEGFR2.

To further examine the question of localization of VEGFR2 and p-VEGFR2, we stained hRECs with their respective antibodies (**Fig. 4**). Before VEGF-A stimulation, the majority of VEGFR2 was localized near nuclei as in (Manickam et al., 2011; Yamada et al., 2014). And a portion of VEGFR2 was also observed at cell-cell junctions in both control peptide-treated and KAI-treated cells (**Fig. 4A**). Three min after VEGF-A stimulation, phosphorylation of VEGFR2 (Y1175) was mainly observed at cell-cell junctions (**Fig. 4A**). KAI treatment did not alter the initial area where VEGFR2 is phosphorylated at 3 min. In control peptide-treated hRECs, p-VEGFR2 was rapidly internalized and distributed in the cells at 15 min after VEGF-A stimulation. However, p-VEGFR2 accumulated in KAI-treated hRECs at 15 min and 30 min after VEGF-A stimulation. The number of cells showing p-VEGFR2 accumulation was significantly increased in KAI-treated cells at 15 min and 30 min after VEGF-A stimulation (**Fig. 4D**). The fluorescence intensity of p-VEGFR2 and VEGFR2 were not different between the two groups (**Fig. 4B, C**), indicating that a similar amount of p-VEGFR2 was distributed in control and accumulated in KAI-treated hREC. Total VEGFR2 was also largely distributed throughout control cells, whereas total VEGFR2 remained near nuclei in KAI-treated cells at 15 min and 30 min after VEGF-A stimulation (**Fig. 4A**). These results together showed that VEGFR2 is first phosphorylated at cell-cell junction, then phosphorylated VEGFR2 is internalized to induce signaling, which releases of VEGFR2 from Golgi (Yamada et al., 2014). Thus both phosphorylated and unphosphorylated VEGFR2 were distributed throughout the cells in control. KAI treatment inhibited the release of unphosphorylated VEGFR2 from Golgi, consistent with the effects of knockdown of *KIF13B* (Yamada et al., 2014). The distribution of p-VEGFR2 was also inhibited by KAI treatment, resulting in accumulation of p-VEGFR2. Phosphorylated VEGFR2 is either dephosphorylated and recycled to the cell surface or undergoes degradation (Simons et al., 2016). Thus, the question arises about the disposition of VEGFR2 during the inhibition of VEGFR2 trafficking.

**Fig. 4.**
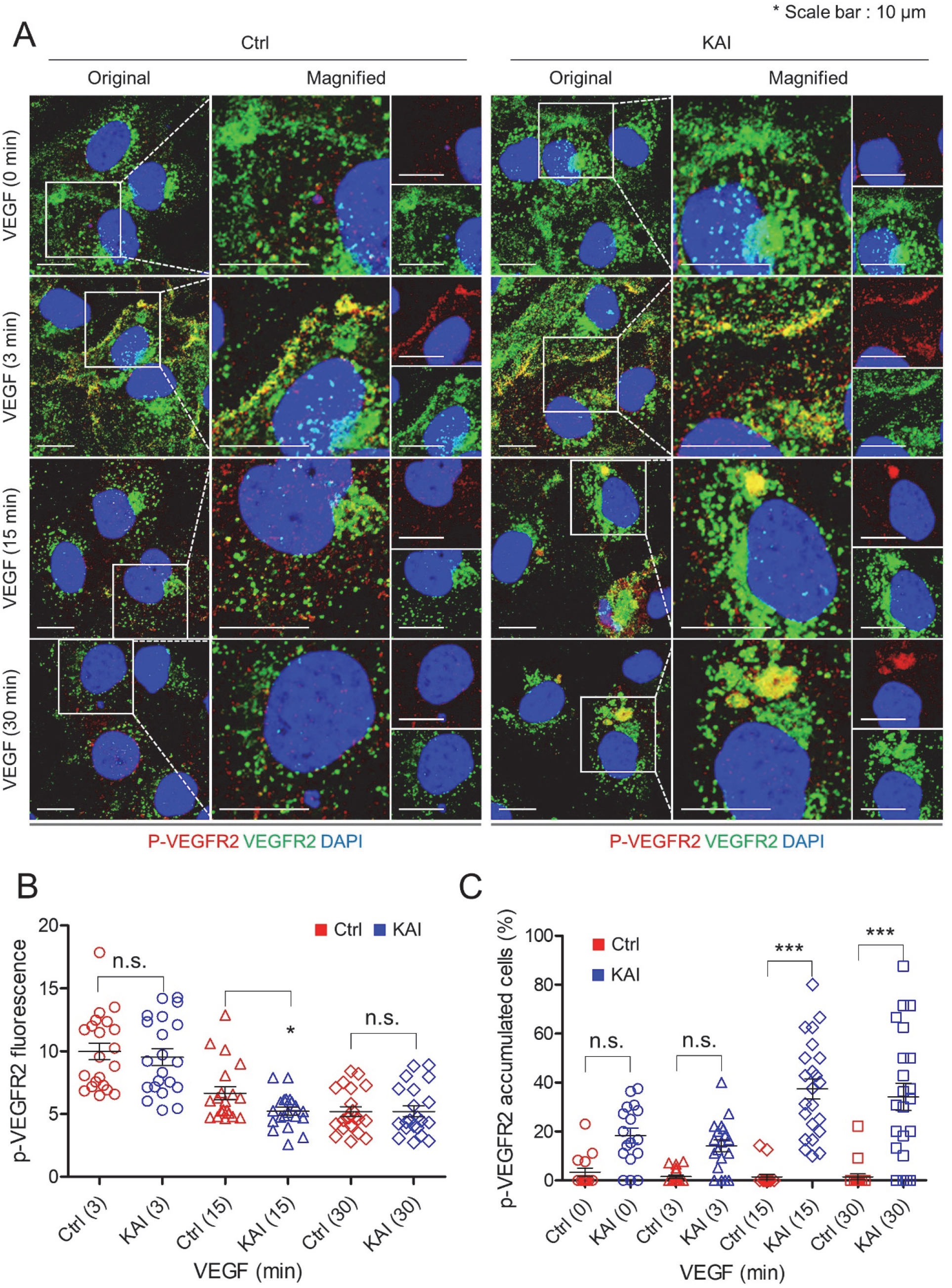
KIF13B regulates endosomal trafficking of phosphorylated VEGFR2. **(A, B, C, D)** Immunostaining for p-VEGFR2 (Y1175, red) and total VEGFR2 (green) in hREC pretreated with either control peptide or KAI (10 µM), and stimulated with VEGF-A (50 ng/mL) for indicated time periods. Scale bars; 10 µm. The intensity of p-VEGFR2 was quantified by Image J and shown as mean ± SE in graph B. N=23, 22, 23, 24, 23, 23, 24 and 24 for pictures in ctrl (0), ctrl (3), ctrl (15), ctrl (30), KAI (0), KAI (3), KAI (15) and KAI (30) were analyzed from 4 independent experiments. Unpaired t-test, * indicates p<0.05, ** indicates p<0.01 and n.s. indicates p>0.05. The number of cells with p-VEGFR2 accumulation (accumulated particles bigger than 5 µm^2^) among the total number of the cells was counted and shown as mean ± SE in graph C. N=15, 20, 18, 21, 17, 19, 23 and 22 for pictures in ctrl (0), ctrl (3), ctrl (15), ctrl (30), KAI (0), KAI (3), KAI (15) and KAI (30) were analyzed from 4 independent experiments. One-way ANOVA with post hoc multiple comparisons. *** indicates p<0.001 and n.s. indicates p>0.05.

### Inhibition of VEGFR2 recycling localizes VEGFR2 in late endosomes

Rab family GTPases localize to specific intracellular compartments where they regulate cargo trafficking (Stenmark, 2009). Internalized VEGFR2 localizes at Rab5A positive early endosome, then post-dephosphorylation, VEGFR2 enters either Rab11-mediated recycling pathway or Rab7-mediated degradation pathway. To determine how KIF13B contributes to VEGFR2 trafficking, we performed co-immunoprecipitation with Rab family proteins, Rab5A, Rab11, and Rab7, and analyzed binding proteins by western blotting (**Fig. 5**). In scrambled peptide-treated control hRECs, VEGFR2 and KIF13B were co-immunoprecipitated with Rab5A (**Fig. 5A**). KAI treatment decreased KIF13B on Rab5A-positive vesicles, whereas VEGFR2 remained co-precipitated with Rab5A (**Fig. 5A, B**). KAI treatment reduced VEGFR2 associated with Rab11-positive recycling vesicle (**Fig 5C, D**), suggesting reduced recycling in KAI-treated cells. The result consistent with the biotinylation assay (**Fig. 3**) shows the crucial role of KIF13B in mediating recycling of internalized VEGFR2. Upon inhibition of VEGFR2 trafficking by KAI treatment, we observed accumulation of p-VEGFR2 near nuclei (**Fig. 4A**). Inhibition of VEGFR2 recycling may thus traffic it to the degradation pathway.

**Fig. 5.**
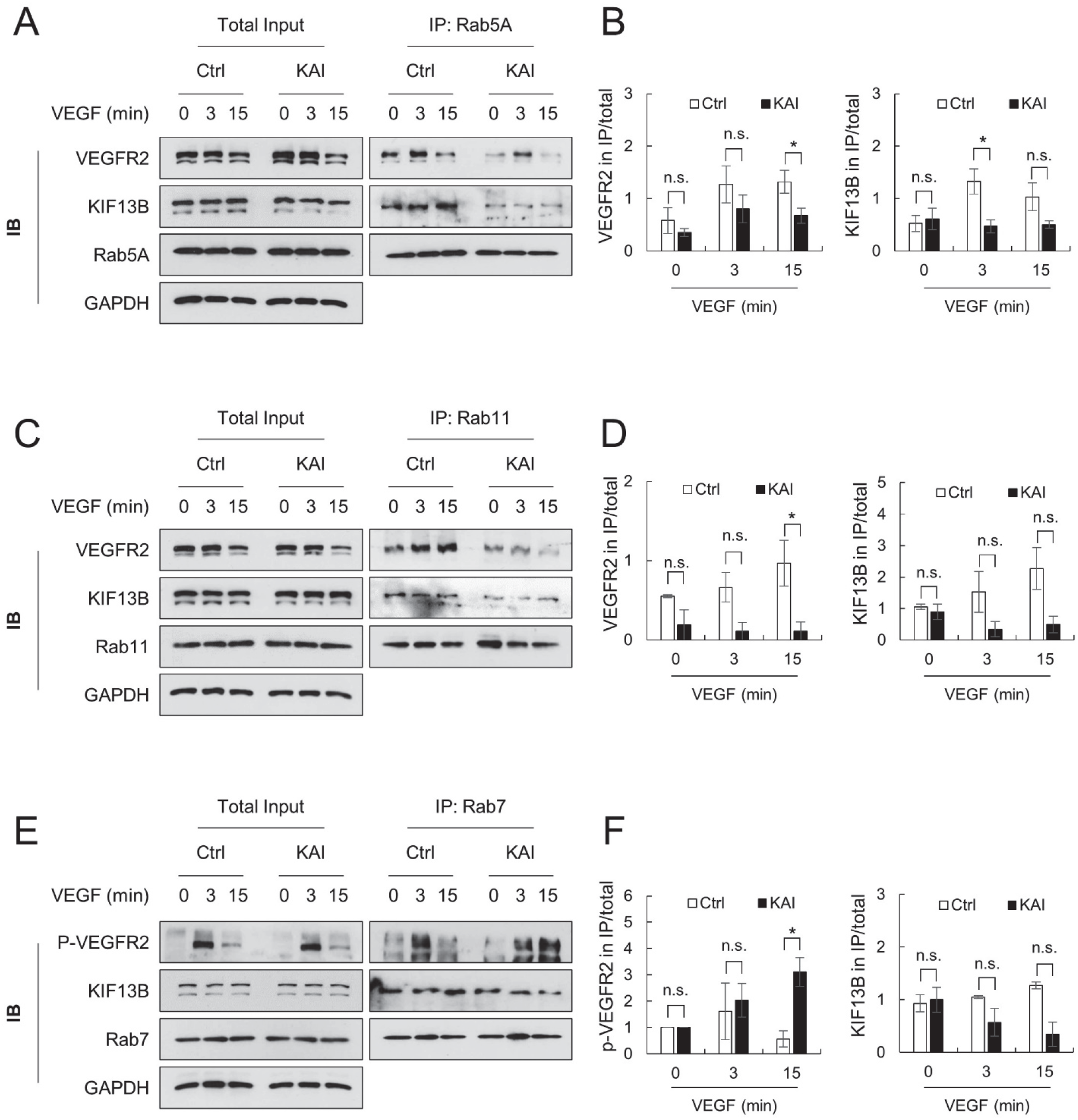
Association of VEGFR2 to Rab family proteins is impaired by KAI, an inhibitor for KIF13B-mediated VEGFR2 trafficking. **(A, B)** Association of VEGFR2 and KIF13B with Rab5A-positive early endosome in hREC pretreated with either scrambled control peptide or KAI (10 µM). After VEGF-A stimulation (50 ng/mL) for indicated time periods, proteins were co-immunoprecipitated (IP) with Rab5A and analyzed by western blotting. Representative blots were shown in A. Quantification of the proteins co-immunoprecipitated with Rab5A was normalized by proteins in the total lysate and shown as mean ± SE in the graph B. N=4. T-test was performed for statistical analysis in ctrl and KAI groups with the same time point of VEGF stimulation. * indicates p<0.05. n.s. indicates p>0.05. **(C, D)** Association of VEGFR2 and KIF13B with Rab11-positive recycling vesicles in hREC pretreated with either scrambled control peptide or KAI (10 µM). After VEGF-A stimulation (50 ng/mL) for indicated time points, proteins were co-immunoprecipitated with Rab11 and analyzed by western blotting. Representative blots were shown in C. Quantification of the proteins co-immunoprecipitated with Rab11 was normalized by proteins in the total lysate and shown as mean ± SE in the graph D. N=4. T-test was performed for statistical analysis in ctrl and KAI groups with the same time point of VEGF stimulation. * indicates p<0.05. n.s. indicates p>0.05. **(E, F)** Association of phosphorylated VEGFR2 (Y1175), and KIF13B with Rab7-positive late endosome in hREC pretreated with either scrambled control peptide or KAI (10 µM). After VEGF-A stimulation (50 ng/mL) for indicated time points, proteins were co-immunoprecipitated with Rab7, and analyzed by western blotting. Representative blots were shown in E. Quantification of the proteins co-immunoprecipitated with Rab7 was normalized by proteins in the total lysate and shown as mean ± SE in the graph F. N=4. T-test was performed for statistical analysis in ctrl and KAI groups with the same time point of VEGF stimulation. * indicates p<0.05. n.s. indicates p>0.05.

To test address this concept, we performed co-immunoprecipitation with a late endosome marker, Rab7 (**Fig. 5E, F**). In control hRECs, p-VEGFR2 (Y1175) was detected in Rab7-positive late endosome at 3 min after VEGF-A stimulation and dephosphorylated at 15 min (**Fig. 5E**). Interestingly, in KAI-treated hREC, accumulation of p-VEGFR2 was observed in Rab7-positive late endosome (**Fig. 5E**).

To confirm the western blotting results, we stained hREC with Rab11 and VEGFR2 (**Fig. 6**). In control peptide-treated cells, colocalization of VEGFR2 with recycling vesicle marker Rab11 was observed, but KAI treatment significantly reduced colocalization at 3 min and 15 min after VEGF-A stimulation (**Fig. 6A, B**). In contrast, p-VEGFR2 (Y1175) was highly colocalized with the late endosome marker Rab7 in KAI-treated cells (**Fig. 7**). At 3 min after VEGF-A stimulation, phosphorylation of VEGFR2 was observed at cell-cell junctions in both control and KAI-treated hREC, consistent with **Fig. 4**. KAI treatment induced accumulation of p-VEGFR2, which was colocalized with Rab7 (**Fig. 7A, B**). We further analyzed colocalization of p-VEGFR2 (Y1175) and lysosome marker LAMP2 (**Fig. 8**). KAI induced accumulation of p-VEGFR2 in LAMP2 positive lysosomes at 15 min and 30 min after VEGF-A stimulation, compared with control (**Fig. 8A, B**). Together, these data show that inhibition of VEGFR2 trafficking by KAI treatment traffics VEGFR2 to late endosome and lysosome for degradation.

**Fig. 6.**
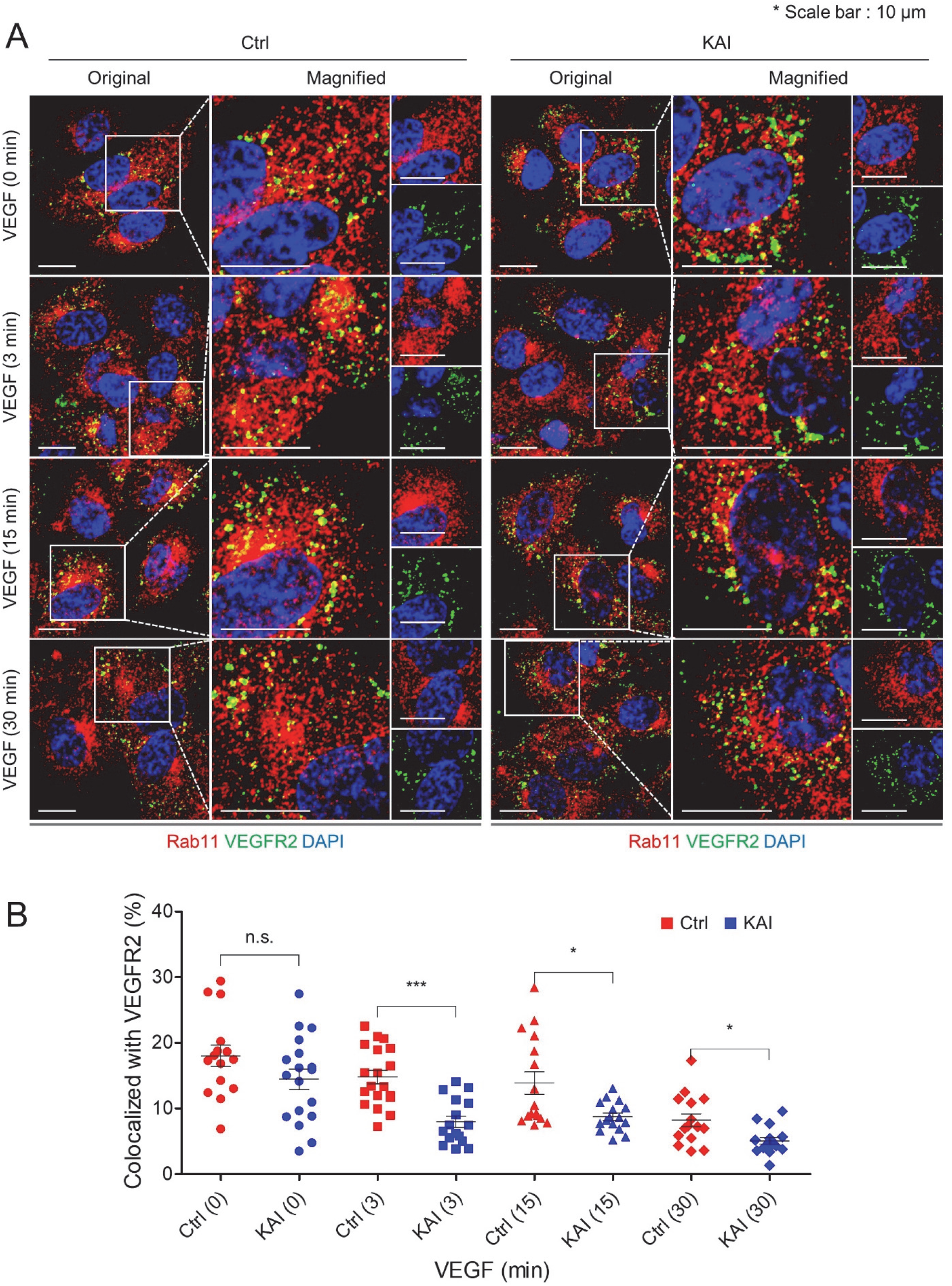
Inhibition of KIF13B reduced association of VEGFR2 to recycling vesicle. **(A, B, C, D)** Immunostaining for VEGFR2 (green) and Rab11 (red) in hREC pretreated with either control peptide or KAI (10 µM), and stimulated with VEGF-A (50 ng/mL) for indicated time periods. Scale bars; 10 µm. The colocalization of VEGFR2 and Rab11 was quantified by Image J and shown as mean ± SE in graph B. N=15, 18, 19, 16, 16, 16, 15 and 16 for pictures in ctrl (0), ctrl (3), ctrl (15), ctrl (30), KAI (0), KAI (3), KAI (15) and KAI (30) were analyzed from 3 independent experiments. One-way ANOVA with post hoc multiple comparisons. * indicates p<0.05, ** indicates p<0.01 and n.s. indicates p>0.05.

**Fig. 7.**
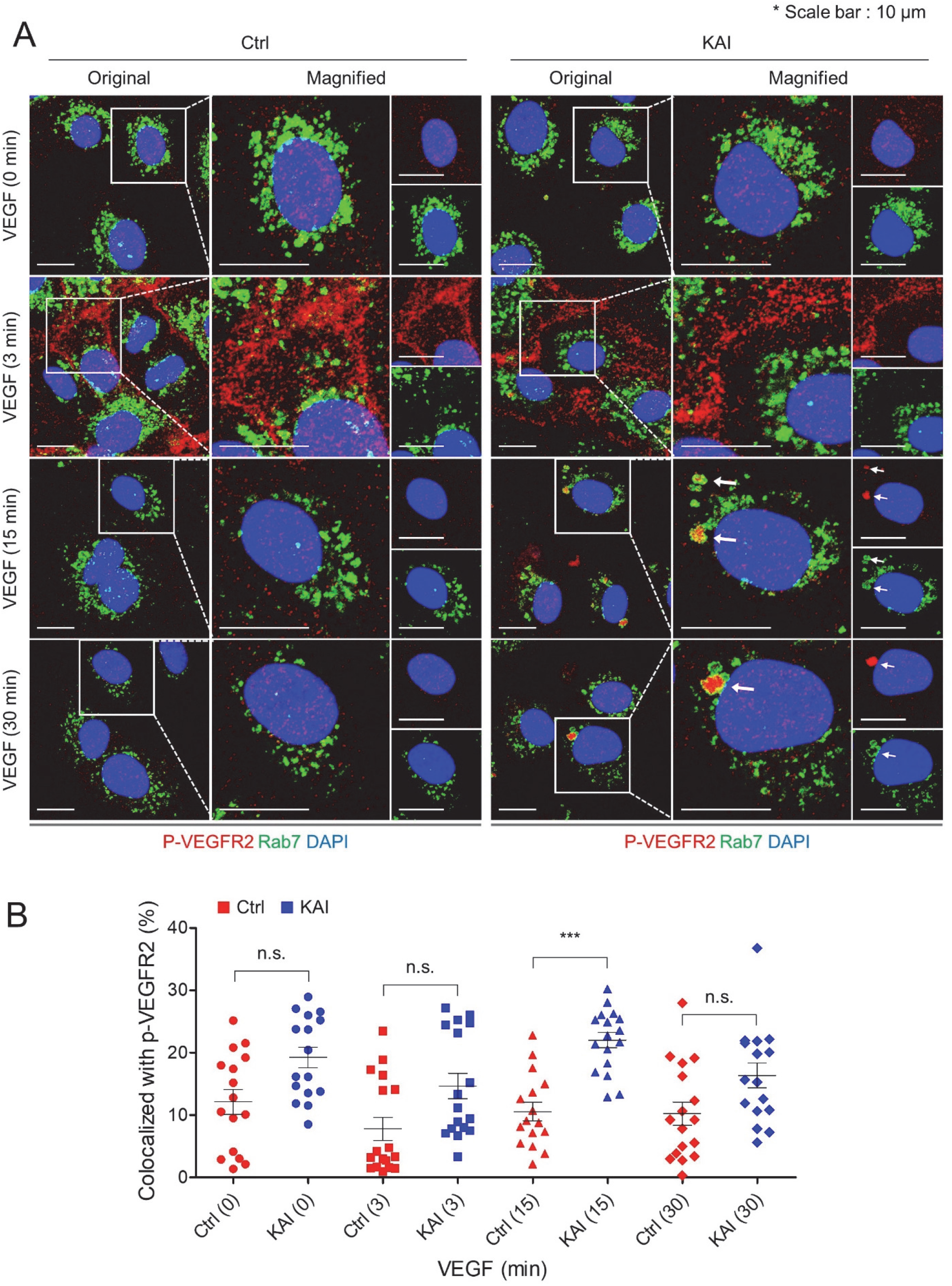
Failure of VEGFR2 trafficking sends p-VEGFR2 to Rab7-positive late-endosome. Immunostaining for p-VEGFR2 (Y1175, red) and Rab7 (green) in hREC pretreated with either control peptide or KAI (10 µM), and stimulated with VEGF-A (50 ng/mL) for indicated time periods. Scale bars; 10 µm. The colocalization of p-VEGFR2 and Rab7 was quantified by Image J and shown as mean ± SE in graph B. N=16, 16, 17, 17, 16, 17, 17 and 16 for pictures in ctrl (0), ctrl (3), ctrl (15), ctrl (30), KAI (0), KAI (3), KAI (15) and KAI (30) were analyzed from 3 independent experiments. One-way ANOVA with post hoc multiple comparisons. *** indicates p<0.001 and n.s. indicates p>0.05.

**Fig. 8.**
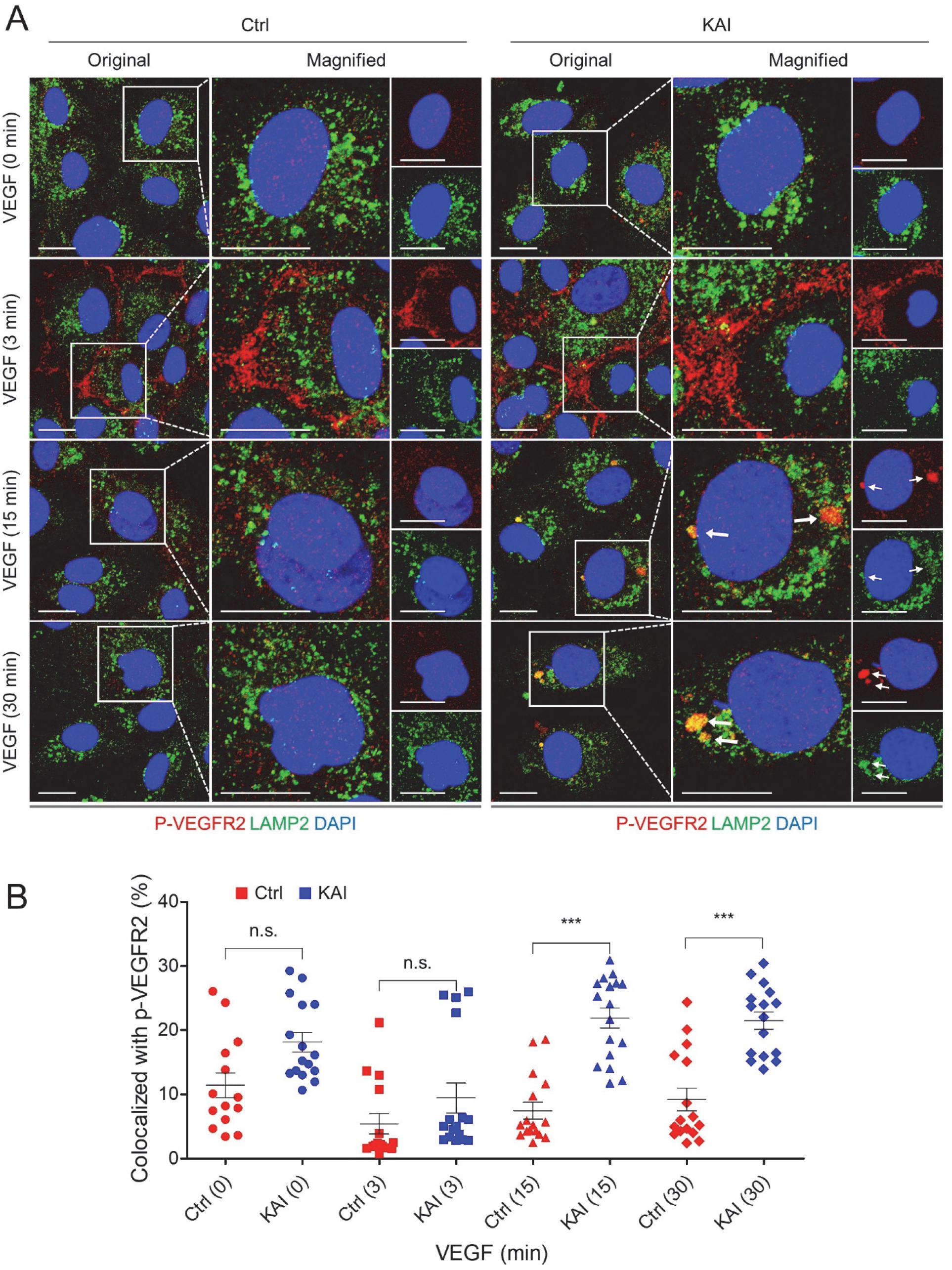
Failure of VEGFR2 trafficking sends p-VEGFR2 to lysosome. **(A)** Immunostaining for p-VEGFR2 (Y1175, red) and LAMP2 (green) in hREC pretreated with either control peptide or KAI (10 µM), and stimulated with VEGF-A (50 ng/mL) for indicated time periods. Scale bars; 10 µm. **(B)** The colocalization of p-VEGFR2 and LAMP2 was quantified by Image J and shown as mean ± SE in graph B. N=14, 16, 15, 16, 16, 17, 16 and 16 for pictures in ctrl (0), ctrl (3), ctrl (15), ctrl (30), KAI (0), KAI (3), KAI (15) and KAI (30) were analyzed from 3 independent experiments. One-way ANOVA with post hoc multiple comparisons. *** indicates p<0.001 and n.s. indicates p>0.05.

We also quantified the total amount of VEGFR2 after VEGF-A stimulation in control peptide-treated hREC and KAI-treated hREC (**Fig. 9A, B**). Compared to control, KAI treatment reduced the total amount of VEGFR2 at 15 min after VEGF-A stimulation (**Fig. 9A, B**).

**Fig. 9.**
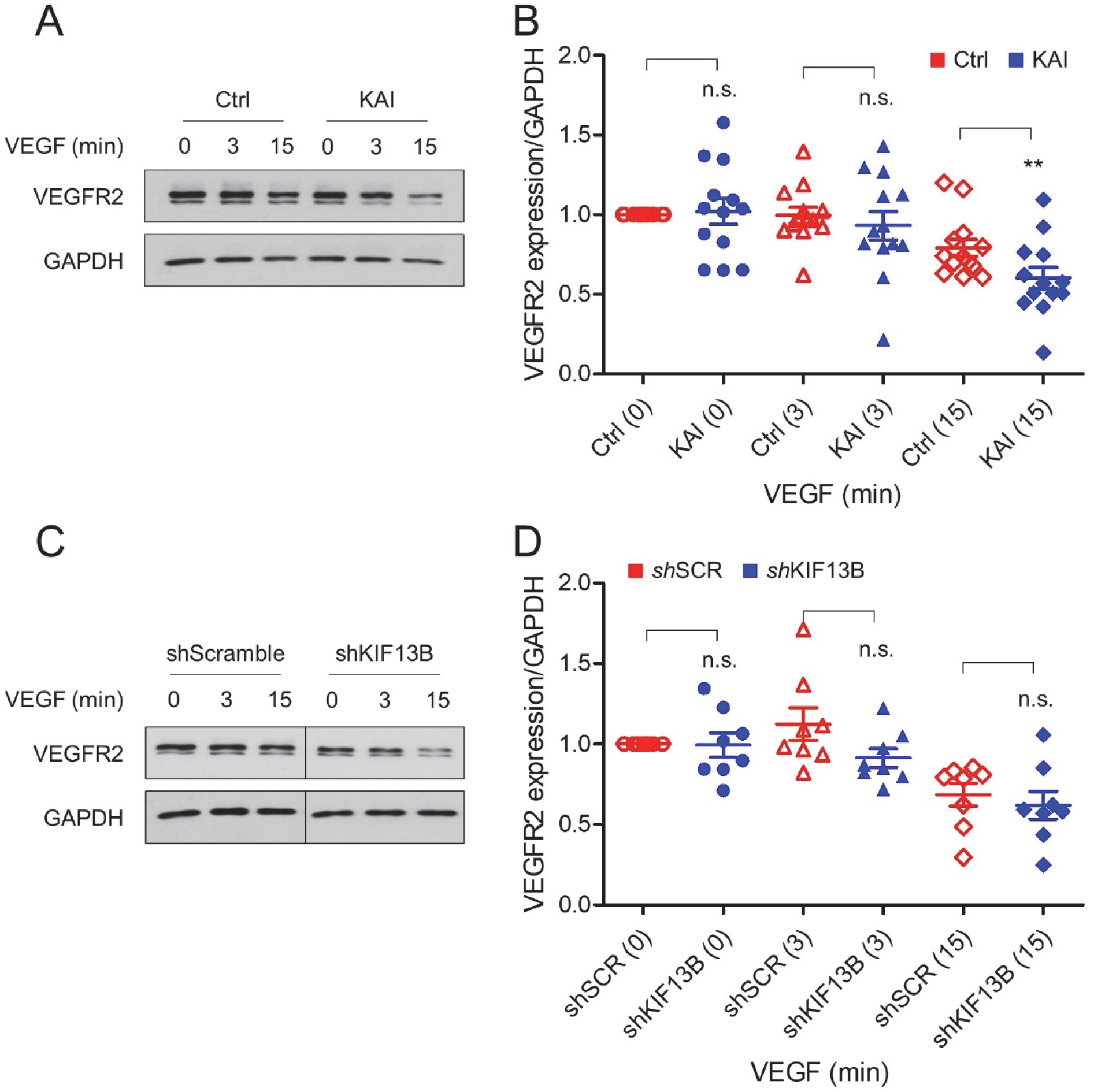
Inhibition of KIF13B function targets VEGFR2 for degradation. **(A, B)** Degradation of VEGFR2 after VEGF-A stimulation (50 ng/mL) in hREC treated with either scrambled control peptide or KAI peptide (10 µM). Representative blots were shown in A. Quantification of VEGFR2 was normalized by loading control GAPDH, and shown in graph B as mean ± SE. N=13. T-test was performed for statistical analysis in ctrl and KAI groups with the same time point of VEGF stimulation. ** indicates p<0.01. **(C, D)** Degradation of VEGFR2 after VEGF-A stimulation (50 ng/mL) in hREC transduced with scrambled shRNA or shRNA-*KIF13B*. Representative blots were shown in C. Intensity of the bands of VEGFR2 was normalized by loading control GAPDH, and shown in graph D as mean ± SE. N=8. T-test was performed for statistical analysis in scrambled shRNA and shRNA-*KIF13B* groups with the same time point of VEGF stimulation. N.s. indicates no significant difference.

Degradation of VEGFR2 was also tested in scrambled shRNA-transduced control and shRNA-*KIF13B*-transduced cells (**Fig. 9C, D**). Knockdown of *KIF13B* tended to decrease the total amount of VEGFR2, although it was not statistically significant (**Fig. 9C, D**). These results together show that KIF13B contributes recycling of internalized VEGFR2. Inhibition of KIF13B traffics VEGFR2 to lysosomal degradation instead of recycling and limits the amount of VEGFR2 on the cell surface. Overall, the results show that VEGFR2 recycling is essential for VEGF-A signaling and endothelial permeability.

### Inhibition of VEGFR2 trafficking ameliorates pathological vascular leakage

We examined whether inhibition of VEGFR2 trafficking is an effective strategy to inhibit vascular leakage in wet AMD. We tested the efficacy of KAI on laser-induced vascular leakage in the mouse model (**Fig. 10**). C57BL/6 mice received laser burns on Bruch’s membrane, which mimics disruption of Bruch’s membrane in wet AMD, and induces vascular leakage and neovascularization (Fantin et al., 2017; Gong et al., 2015). Mice were treated with eyedrop of either control peptide or KAI (5 µg/eye in 5 µl in PBS) once a day for 3 days. Three days after laser burns, Evans blue was injected i.p. to examine the Evans blue extravasation on the following day. The control peptide-treated mice showed extravasation of Evans blue at the site with laser burn (**Fig. 10A top panel**). KAI treatment reduced the extravasation of Evans blue (**Fig. 10A bottom panel and 10B**). Laser-induced neovascularization at day 4 was also analyzed by staining with isolectin B4 (ILB4), an endothelial marker. Compared to the control mice, laser-induced neovascularization was also reduced by daily treatment with KAI, but it was not statistically significant (**Fig. 10A, C**). Note, neovascularization at the late stage was significantly reduced by continuous KAI treatment for 14 days (Waters et al., 2021b). As laser-induced vascular leakage also facilitated macrophages recruitment (Fantin et al., 2017), we observed macrophages by staining with marker CD68 (**Fig. 10A, D**). Thus, it appears that KAI treatment also inhibited macrophage recruitment.

**Fig. 10.**
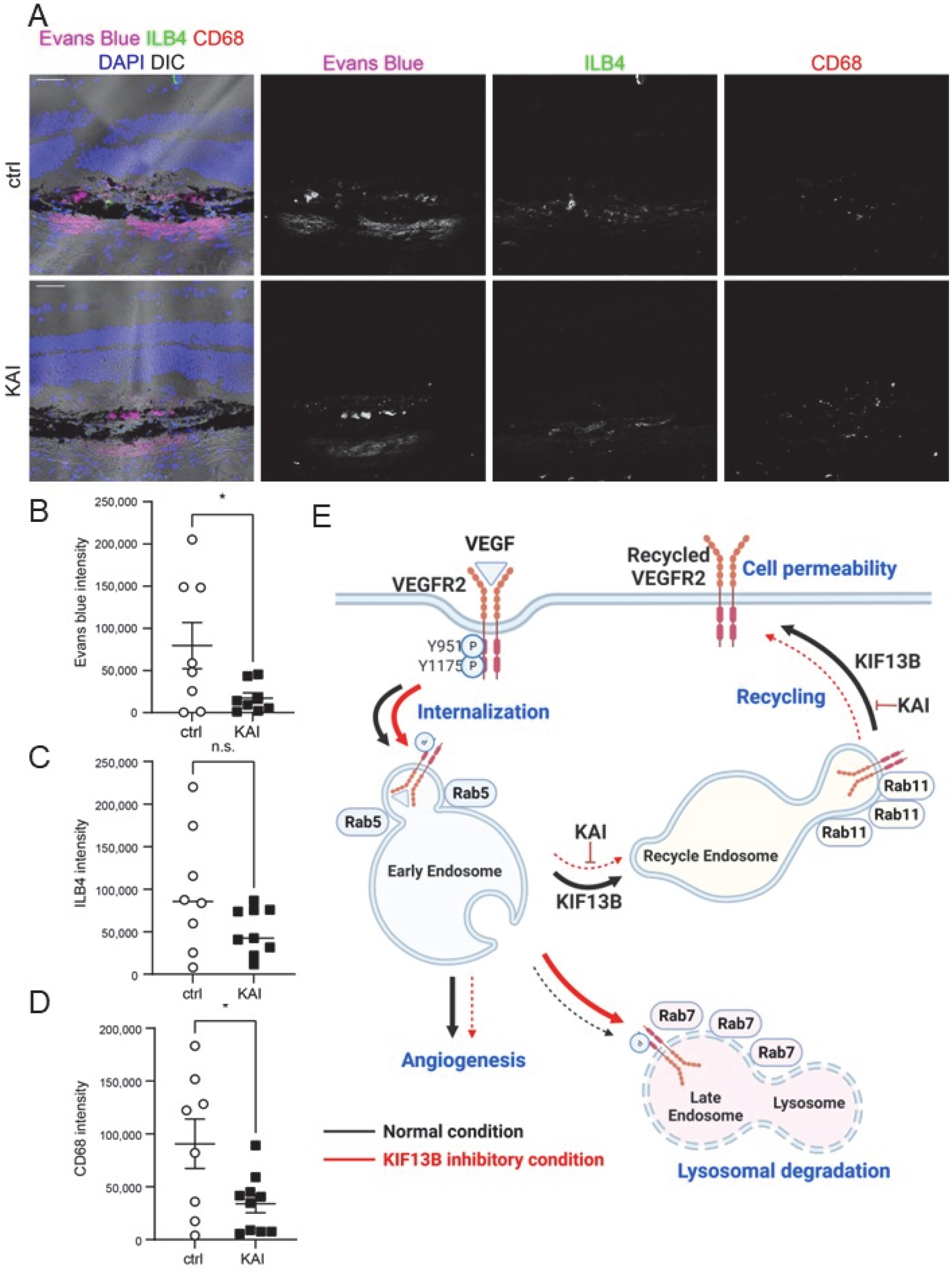
Pharmacological inhibition of VEGFR2 trafficking ameliorates vascular leakage in mice. **(A, B, C, D)** Effects of KAI eyedrop on the pathological vascular leakage in mice. After receiving laser burns, mice were treated either control peptide (ctrl) or KAI (5 µg/eye in 5 µl PBS) daily. On day 3, Evans blue was injected i.p., and Evans blue leakage was measured 24 hr after injection. Cryosections of the damaged area were also stained with ILB4 and CD68 to visualize neovascularization and macrophage recruitment, respectively. Representative images were shown in A. Scale bars; 50 µm. The intensity of the Evans blue extravasation, ILB4, and CD68 was quantified and shown in the graph as mean ± SE in B, C, D. N=8 and 10 for ctrl and KAI, respectively. **(E)** Schematic showing the working model in this study. VEGF induces internalization of phosphorylated VEGFR2 to Rab5-positive early endosomes. Dephosphorylated VEGFR2 is transferred to Rab11-positive recycling vesicle by KIF13B and further trafficked to the cell surface. When KIF13B function is blocked by either knockdown or pharmacological inhibitor KAI, VEGFR2 trafficking is failed, and phosphorylated VEGFR2 is sent from early endosomes to late endosome and lysosome for degradation. Recycling of VEGFR2 is required for VEGF-induced signaling for endothelial permeability. Thus this pathway can be a therapeutic target for vascular leakage-related diseases such as wet AMD.

## Discussion

Herein we showed that VEGF-A stimulation first induced phosphorylation of VEGFR2 at cell-cell junction. Internalized p-VEGFR2 then trafficked to Rab5 positive early endosome to induce signaling (Nakayama et al., 2013; Sawamiphak et al., 2010; Lanahan et al., 2010). Our data show that KIF13B interacted with internalized VEGFR2 at Rab5-positive early endosome to escort VEGFR2 to the recycling pathway (see **Fig. 10E**). Blocking the interaction between VEGFR2 and KIF13B prevented VEGFR2 dephosphorylation, a necessary step for the receptor to enter the Rab11 recycling pathway (Ballmer-Hofer et al., 2011; Lanahan et al., 2014). Thus, inhibition of VEGFR2 trafficking caused aberrant accumulation of phosphorylated VEGFR2 in lysosomes to be degraded. This recycling of VEGFR2 to the cell surface may be important in tuning VEGF signaling and regulating endothelial permeability, as both KAI treatment and knockdown of *KIF13B* inhibited VEGF-induced endothelial permeability.

Endocytosis of phosphorylated VEGFR2 is mediated by ephrin-B2, PAR-3, Dab2, and aPKC (Nakayama et al., 2013; Sawamiphak et al., 2010). Internalized VEGFR2 is then transported to early endosomes by myosin family molecular motor MYO6 and synectin (Lanahan et al., 2010). In contrast, VEGFR2 trafficking to the cell surface is mediated by KIF13B (Yamada et al., 2014), t-SNARE syntaxin 6 (Manickam et al., 2011), SUMO endopeptidase SENP1 (Zhou et al., 2018), and another myosin family molecular motor MYO1C (Tiwari et al., 2013). The potential interaction and cooperation of KIF13B with these molecules needs further investigation. Nonetheless, we demonstrated the requisite role of KIF13B in VEGFR2 trafficking during recycling, in addition to the known role of KIF13B trafficking of newly synthesized VEGFR2 from the Golgi apparatus (Yamada et al., 2014).

Failure of trafficking misdirect VEGFR2 to the degradation pathway (Yamada et al., 2014; Tiwari et al., 2013; Manickam et al., 2011; Ash et al., 2021; Kofler et al., 2018). Thus, KAI peptide decreased VEGFR2 in early endosomes and recycling endosomes and increased VEGFR2 in Rab7-positive late endosomes at 15 min after VEGF-A stimulation. To keep functional VEGFR2 in the cycle, unfunctional VEGFR2 needs to be rapidly and efficiently removed to avoid disruptive signaling. Ubiquitin isopeptidase USP8 (Smith et al., 2016) and ER-resident ubiquitin E3 ligase RNF121 mediate degradation of immature VEGFR2 (Maghsoudlou et al., 2016). How failed trafficking of VEGFR2 induces its ubiquitination and subsequent degradation remains an open question.

The interaction between Rab family proteins and KIF13B or its homolog has been reported (Huckaba et al., 2011; Bentley et al., 2015; Delevoye et al., 2014). KIF13B interacts with the early endosome through its tail domain (832-1826 aa) (Bentley et al., 2015), which includes the VEGFR2 binding domain (1238-1260 aa) and Cap-gly domain (1703-1768 aa). As internalized VEGFR2 enters Rab5-positive early endosomes (Nakayama et al., 2013; Sawamiphak et al., 2010; Lanahan et al., 2010), KIF13B may interact with early endosomes via VEGFR2. Alternatively, KIF13B may directly interact with Rab5 through its Cap-gly domain, as in the case of kinesin-73, *Drosophila* homolog of KIF13B (Huckaba et al., 2011). As KAI reduced the association of KIF13B to Rab5-positive vesicles, the interaction of KIF13B to early endosome likely occurs via VEGFR2. KIF13A, the closest homolog of KIF13B, interacts with the active form of Rab11 via stalk (351-1307 aa) and tail (1307-1770 aa) (Delevoye et al., 2014). Similarly, KIF13B may interact with Rab11. Alternatively, KIF13B and KIF13A may form a heterodimer (Mills et al., 2019) for the recycling of VEGFR2. It is noteworthy that KIF13B did not interact with the lysosome (Bentley et al., 2015). Together, our data show that KIF13B interacts with internalized VEGFR2 in early endosome and escorts VEGFR2 to the Rab11-mediated recycling pathway. Thus, inhibition of the function of KIF13B prevents recycling of VEGFR2 and leads to lysosomal degradation of VEGFR2.

Our previous studies showed that KIF13B-mediated VEGFR2 trafficking regulated EC migration and sprouting angiogenesis (Yamada et al., 2014) and VEGF-induced vascular leakage over periods of up to weeks (Waters et al., 2021a). The present results, however, show a rapid response to VEGF-A occurring within 3 min. As shown in the present study, the interaction between VEGFR2 in the early endosomes with molecular motors KIF13B and MYO1C restore VEGFR2 at the cell surface within minutes. Kinesins transport their cargo for 50-200 mm per day (Hirokawa et al., 2009), that is, 35-140 µm/min. The speed of kinesins is sufficiently rapid to transport cargo to the cell surface within minutes in endothelial cells (50-70 µm diameter). Our data demonstrated that inhibition of KIF13B-mediated VEGFR2 trafficking impaired VEGF signaling within the same minutes time frame.

Then, how is VEGF signaling regulated by VEGFR2 trafficking? Abrogation of KIF13B-mediated VEGFR2 trafficking reduced phosphorylation of VEGFR2 greater degree at Y951 than Y1175 sites, even though the initial cell-surface VEGFR2 levels were not altered. VEGF stimulation simultaneously induces auto-phosphorylation of VEGFR2 at multiple tyrosine residues, Y951, Y1054/1059, Y1175, and Y1214 (Corti and Simons, 2017). Each phosphorylation site induces unique downstream signaling pathways. Phosphorylation at Y1175 regulates cell migration, proliferation and homeostasis, whereas phosphorylation at Y951 regulates permeability and survival (Simons et al., 2016). Dephosphorylation of VEGFR2 is mediated by multiple phosphatases with overlapping and preferred target residues. Cell membrane phosphatase vascular endothelial protein Tyr phosphatases (VE-PTPs) dephosphorylates both Y951 and Y1175 of VEGFR2 (Corti and Simons, 2017). Intracellular phosphatases Src homology-2 domain-containing phosphatase 1 (Shp1) dephosphorylates Y1175 and Y1054/59, but not Y951 (Corti and Simons, 2017). In contrast, Shp2 dephosphorylates VEGFR2 at Y951, Y996, and Y1059, but not Y1175 (Sinha et al., 2009). Regulation of phosphorylation status of VEGFR2 by its trafficking has been demonstrated (Lanahan et al., 2010; Chittenden et al., 2006; Kofler et al., 2018; Lanahan et al., 2013). When trafficking of internalized VEGFR2 from periphery to early endosome is inhibited, VEGFR2 can be dephosphorylated by membrane phosphatases (Lanahan et al., 2013, 2010). Similarly, when recycling of internalized VEGFR2 is inhibited, VEGFR2 can be dephosphorylated by intracellular phosphatases (Kofler et al., 2018). In this study, we showed reduced phosphorylation of Y951 by inhibition of KIF13B function mediating VEGFR2 trafficking, whereas phosphorylation of Y1175 was sustained. Whether failure of VEGFR2 recycling causes exposure of VEGFR2 to phosphatase, which selectively dephosphorylate VEGFR2 at Y951, is still an open question. An alternative possibility is failure of VEGFR2 recycling reduces the amount of VEGFR2 at the cell surface to be ligated with VEGF. Impaired VEGFR2 trafficking caused accumulation and degradation of VEGFR2 in lysosomes. Thus, VEGF signaling was reduced and release of VEGFR2 from the perinuclear pool was inhibited. Both inhibition of VEGFR2 trafficking from the recycling pool and the newly synthesized pool prevented VEGF signaling, thereby inhibiting downstream signaling mediating endothelial permeability. Thus, targeting VEGFR2 trafficking is potentially a promising anti-endothelial permeability strategy.

## Materials and Methods

### Antibodies and reagents

Rabbit antibodies against VEGFR2, p-VEGFR2 (Y1175), p-VEGFR2 (Y951), Src, p-Src (Y416), Rab11, CD68 were from Cell Signaling Technology (Danvers, MA, USA). Rabbit antibodies against p-VE-cadherin (Y658) (Invitrogen, Carlsbad, CA, USA), p-VE-cadherin (Y685) (Abcam, Cambridge, MA, USA), VE-cadherin (Abcam), GAPDH (Sigma-Aldrich, St. Louis, MO, USA) were also used. The goat antibody against VEGFR2 was from R&D Systems (Minneapolis, MN). Mouse antibodies against Rab5A, Rab7 (Cell Signaling), and LAMP2 (BD Bioscience, Franklin Lakes, NJ, USA) were also used. Mouse monoclonal antibody against KIF13B was kindly provided by Dr. Toshihiko Hanada (Horiguchi et al., 2006; Mills et al., 2019) (Tufts University, Boston, USA). Other reagents used were ILB4 (Vector Laboratories, Burlingame, CA, USA), Alexa 594 anti-rabbit antibody, Alexa 488 anti-mouse antibody, Alexa 488 anti-goat antibody, Alexa 488-streptavidin (Invitrogen), and HRP-conjugated secondary antibodies (Jackson Immunoresearch Laboratories, West Grove, PA, USA). Recombinant human VEGF-A^165^ was purchased from Peprotech (Cranbury, NJ, USA)

### Cell culture

Human retinal endothelial cells (hRECs) (Cell Systems, Kirkland, WA, USA) were maintained with EGM2 MV supplemented with 5% FBS (Lonza Group, Basel, Switzerland).

### Plasmids and lentivirus preparation

Lentiviruses encoding shRNA for human *KIF13B* and scrambled shRNA were as described (Yamada et al., 2014). Titer of the lentiviruses was measured with a lentivirus titration kit (Takara Bio USA, San Jose, CA, USA). The same multiplicity of infection (MOI) of lentiviruses encoding scrambled shRNA and shRNA-*KIF13B* (MOI 100) were used for hREC. 72 hr after infection, cells were used for experiments.

### Peptides

Biotin-KAI and biotin-ctrl were as described (Yamada et al., 2017; Waters et al., 2021b) Scrambled KAI was also designed (TFLTLRDRRSHLPGVSQVPIEVV) and synthesized by a custom synthesized service (Thermo Fisher Scientific, Waltham, MA, USA) and used as control peptide.

### Transendothelial electrical resistance (TEER) assay

To examine the function of KIF13B to regulate VEGF-induced endothelial barrier relaxation, transendothelial electrical resistance (TEER) was measured by electrical cell-substrate impedance sensing (ECIS, Applied BioPhysis inc, Troy, NY). Briefly, hRECs were infected with lentiviruses encoding shRNA-*KIF13B* or scrambled shRNA, and seeded in the 0.1% gelatin-coated electrode chamber (8W10E+C) as 4×10^4^ cells/well. TEER was measured every 30 seconds overnight (4,000 Hz of frequency) to monitor barrier relaxation by VEGF-A (50 ng/mL). Resistance was normalized by the resistance at time zero (when VEGF-A was added) and shown in the graphs.

To examine the effect of KAI treatment, confluent hRECs in the electrode chamber were pretreated with either control peptide or KAI peptide (10 µM) for 2 hr in the complete culture medium. The change of TEER by VEGF-A (50 ng/mL) was measured as described above.

### Western blotting analysis

Confluent hRECs were treated with either control peptide (ctrl) or KAI (10 μM) for 2 hr in the serum starvation media. After stimulation with 50 ng/mL of VEGF-A for indicated time points, cells were lysed by RIPA lysis buffer (Cell Signaling) supplemented with 10 mM sodium fluoride (Sigma), 1 mM sodium orthovanadate (Sigma), 1 mM beta-glycerophosphate (Sigma), and protease inhibitor cocktail (Sigma). Protein concentrations were measured by BCA protein assay kit (ThermoFisher), equal amounts of proteins were analyzed for western blotting. The intensity of the bands was quantified with Image Studio Lite software (LI-COR, Lincoln, NE, USA).

### Immunostaining

Confluent hRECs in 0.1% gelatin-coated EZ slide 4-well chamber (Millipore, Burlington, MA, USA) were pretreated with either scrambled or KAI peptides in serum starvation media for 2 hr. After stimulation with VEGF-A (50 ng/mL) for the indicated time points, cells were fixed with 4% PFA for 20 min at room temperature. Cells were incubated with specific primary antibodies overnight, followed by incubation with fluorescently labeled secondary antibodies with Alexa-594 and Alexa-488 (Invitrogen). Stained cells were observed under a Zeiss confocal LSM 880 META with 63× oil immersion objective lens. The confocal images were all obtained in the same microscope settings, such as gain and scanning time. The intensity was quantified with Image J (NIH).

### Cell surface biotinylation studies

To measure the cell surface pool of VEGFR2, we covalently labeled cell-surface proteins using a membrane-impermeable biotinylation reagent (NHS-SS-biotin; Thermo Fisher) as described previously (Yamada et al., 2014) Total cell proteins and biotinylated surface proteins were analyzed by western blotting using anti-VEGFR2 antibody.

### Biotinylation of internalized VEGFR2

For assessment of the internalized pool of VEGFR2 after VEGF stimulation, biotinylation of cell surface proteins was performed before VEGF-A stimulation as described (Nitzsche et al., 2021). Briefly, after serum starvation for 2 hours, cells were incubated 0.2 mg/mL of sulfo-NHS-SS-biotin for 20 min at 4°C. After biotin labeling on the surface proteins, cells were stimulated with VEGF-A (50 ng/mL) for indicated time points. Cells were washed with cold PBS, and cell surface biotin was cleaved off by incubating the cells on ice with 100 mM of membrane-impermeable reducing agent MESMA (2-mercaptoethane sulphonic acid (Sigma) as described (Nitzsche et al., 2021). Cells were lysed with lysis buffer (10 mM HEPES, 150 mM NaCl, 1% TritonX100, 0.5% NP40, 1 mM EGTA, 1 mM EDTA, 10 mM sodium fluoride, 1 mM sodium orthovanadate, 1 mM beta-glycerophosphate, and protease inhibitor cocktail). Equal amounts of protein lysates were precipitated with streptavidin-agarose beads. After washing the beads extensively with lysis buffer, proteins were analyzed by western blotting.

### Co-immunoprecipitation

After stimulation of the cells with VEGF-A (50 ng/mL) for indicated time points, cells were lysed with lysis buffer as described above. Total protein lysates were precleared with normal IgG conjugated with agarose (Santa Cruz) for 1 hr at 4°C, then proceeded for immunoprecipitation with specific antibodies (Rab5A, Rab 11 and Rab7) and Protein A/G bead (Thermo Fisher) for overnight at 4°C. Proteins co-immunoprecipitated with Rab family antibodies were analyzed by Western blotting.

### Laser-induced vascular leakage

Under anesthesia, C57BL/6 received the laser burns as described (Waters et al., 2021b). Mice received eyedrop of either KAI (5 μg/eye in 5 μL PBS) or ctrl (5 μg/eye) daily on the eyes with laser burns from day 0 to day 3. On day 3 post-lasering, 250 µL of Evans blue (1% in saline) was injected i.p. Eyes were isolated 24 hr later, fixed in 2% paraformaldehyde (PFA) for 30 min and frozen in optimal cutting density compound (OCT) (Sakura Finetek USA, Torrance, CA, USA) in 2 methyl 1 butane in liquid nitrogen. Cryosections of the damaged area were stained with biotinylated ILB4 followed by Alexa 488-streptavidin, or CD68 antibody and Alexa 594-anti-rabbit antibody and counterstained with DAPI. The intensity of Evans blue, ILB4, CD68 was measured using Image J and plotted in the graph using GraphPad Prism 9 (GraphPad, San Diego, CA, USA).

### Statistical analysis

Data were analyzed with GraphPad Prism 9 (GraphPad). Two samples were compared by Student’s *t*-test. More than 2 samples were analyzed by one-way analysis of variance (ANOVA), followed by post hoc comparisons.

## Acknowledgments

We thank Dr. Toshihiko Hanada (Tufts University, Boston, MA, USA) for sharing a monoclonal antibody against KIF13B (Horiguchi et al., 2006). Portions of this work were carried out in the Fluorescent Imaging Core via the Research Resources Center (RRC) at the University of Illinois at Chicago (UIC).

## Author contributions

Conceptualization: HDC, KHY

Methodology: HDC, KHY

Investigation: HDC, CZ, KT, NAS, TN

Visualization: HDC, CZ, KT, NAS

Funding acquisition: HDC, TN, KHY

Project administration: KHY

Supervision: KHY

Writing—original draft: HDC, KHY

Writing—review & editing: KHY

## Data and materials availability

The authors declare that the data supporting the findings of this study are available within the paper and its Supplementary materials. Further information and requests for resources and reagents should be directed to and will be fulfilled by the Lead Contact, Kaori Yamada.

## References

Adam, A.P., A.L. Sharenko, K. Pumiglia, and P.A. Vincent. 2010. Src-induced Tyrosine Phosphorylation of VE-cadherin Is Not Sufficient to Decrease Barrier Function of Endothelial Monolayers* ä. J Biol Chem. 285:7045–7055. doi:10.1074/jbc.m109.079277.

Apte, R.S., D.S. Chen, and N. Ferrara. 2019. VEGF in Signaling and Disease: Beyond Discovery and Development. Cell. 176:1248–1264. doi:10.1016/j.cell.2019.01.021.

Ash, D., V. Sudhahar, S.-W. Youn, M.N. Okur, A. Das, J.P. O’Bryan, M. McMenamin, Y. Hou, J.H. Kaplan, T. Fukai, and M. Ushio-Fukai. 2021. The P-type ATPase transporter ATP7A promotes angiogenesis by limiting autophagic degradation of VEGFR2. Nat Commun. 12:3091. doi:10.1038/s41467-021-23408-1.

Ballmer-Hofer, K., A.E. Andersson, L.E. Ratcliffe, and P. Berger. 2011. Neuropilin-1 promotes VEGFR-2 trafficking through Rab11 vesicles thereby specifying signal output. Blood. 118:816–826. doi:10.1182/blood-2011-01-328773.

Bentley, M., H. Decker, J. Luisi, and G. Banker. 2015. A novel assay reveals preferential binding between Rabs, kinesins, and specific endosomal subpopulations. The Journal of cell biology. 208:273–81. doi:10.1083/jcb.201408056.

Chittenden, T.W., F. Claes, A.A. Lanahan, M. Autiero, R.T. Palac, E.V. Tkachenko, A. Elfenbein, C.R. de Almodovar, E. Dedkov, R. Tomanek, W. Li, M. Westmore, J. Singh, A. Horowitz, M.J. Mulligan-Kehoe, K.L. Moodie, Z.W. Zhuang, P. Carmeliet, and M. Simons. 2006. Selective Regulation of Arterial Branching Morphogenesis by Synectin. Dev Cell. 10:783–795. doi:10.1016/j.devcel.2006.03.012.

Claesson-Welsh, L., E. Dejana, and D.M. McDonald. 2020. Permeability of the Endothelial Barrier: Identifying and Reconciling Controversies. Trends Mol Med. 27:314–331. doi:10.1016/j.molmed.2020.11.006.

Corti, F., and M. Simons. 2017. Modulation of VEGF receptor 2 signaling by protein phosphatases. Pharmacol Res. 115:107–123. doi:10.1016/j.phrs.2016.11.022.

Delevoye, C., S. Miserey-Lenkei, G. Montagnac, F. Gilles-Marsens, P. Paul-Gilloteaux, F. Giordano, F. Waharte, M.S. Marks, B. Goud, and G. Raposo. 2014. Recycling Endosome Tubule Morphogenesis from Sorting Endosomes Requires the Kinesin Motor KIF13A. Cell Reports. 6:445–454. doi:10.1016/j.celrep.2014.01.002.

Duong, C.N., and D. Vestweber. 2020. Mechanisms Ensuring Endothelial Junction Integrity Beyond VE-Cadherin. Front Physiol. 11:519. doi:10.3389/fphys.2020.00519.

Eelen, G., L. Treps, X. Li, and P. Carmeliet. 2020. Basic and Therapeutic Aspects of Angiogenesis Updated. Circ Res. 127:310–329. doi:10.1161/circresaha.120.316851.

Fantin, A., A. Lampropoulou, V. Senatore, J.T. Brash, C. Prahst, C.A. Lange, S.E. Liyanage, C. Raimondi, J.W. Bainbridge, H.G. Augustin, and C. Ruhrberg. 2017. VEGF165-induced vascular permeability requires NRP1 for ABL-mediated SRC family kinase activation. J Exp Medicine. 214:1049–1064. doi:10.1084/jem.20160311.

Gong, Y., J. Li, Y. Sun, Z. Fu, C.H. Liu, L. Evans, K. Tian, N. Saba, T. Fredrick, P. Morss, J. Chen, and L.E. Smith. 2015. Optimization of an Image-Guided Laser-Induced Choroidal Neovascularization Model in Mice. Plos One. 10:e0132643. doi:10.1371/journal.pone.0132643.

Hirokawa, N., Y. Noda, Y. Tanaka, and S. Niwa. 2009. Kinesin superfamily motor proteins and intracellular transport. Nat Rev Mol Cell Biol. 10:682–96. doi:10.1038/nrm2774.

Horiguchi, K., T. Hanada, Y. Fukui, and A.H. Chishti. 2006. Transport of PIP3 by GAKIN, a kinesin-3 family protein, regulates neuronal cell polarity. The Journal of cell biology. 174:425–36. doi:10.1083/jcb.200604031.

Huckaba, T.M., A. Gennerich, J.E. Wilhelm, A.H. Chishti, and R.D. Vale. 2011. Kinesin-73 Is a Processive Motor That Localizes to Rab5-containing Organelles. J Biol Chem. 286:7457–7467. doi:10.1074/jbc.m110.167023.

Kofler, N., F. Corti, F. Rivera-Molina, Y. Deng, D. Toomre, and M. Simons. 2018. The Rab-effector protein RABEP2 regulates endosomal trafficking to mediate vascular endothelial growth factor receptor-2 (VEGFR2)-dependent signaling. J Biol Chem. 293:4805–4817. doi:10.1074/jbc.m117.812172.

Lanahan, A., X. Zhang, A. Fantin, Z. Zhuang, F. Rivera-Molina, K. Speichinger, C. Prahst, J. Zhang, Y. Wang, G. Davis, D. Toomre, C. Ruhrberg, and M. Simons. 2013. The Neuropilin 1 Cytoplasmic Domain Is Required for VEGF-A-Dependent Arteriogenesis. Dev Cell. 25:156–168. doi:10.1016/j.devcel.2013.03.019.

Lanahan, A.A., K. Hermans, F. Claes, J.S. Kerley-Hamilton, Z.W. Zhuang, F.J. Giordano, P. Carmeliet, and M. Simons. 2010. VEGF receptor 2 endocytic trafficking regulates arterial morphogenesis. Dev Cell. 18:713–24. doi:10.1016/j.devcel.2010.02.016.

Lanahan, A.A., D. Lech, A. Dubrac, J. Zhang, Z.W. Zhuang, A. Eichmann, and M. Simons. 2014. PTP1b Is a Physiologic Regulator of Vascular Endothelial Growth Factor Signaling in Endothelial Cells. Circulation. 130:902–909. doi:10.1161/circulationaha.114.009683.

Maghsoudlou, A., R.D. Meyer, K. Rezazadeh, E. Arafa, J. Pudney, E. Hartsough, and N. Rahimi. 2016. RNF121 Inhibits Angiogenic Growth Factor Signaling by Restricting Cell Surface Expression of VEGFR-2. Traffic. 17:289–300. doi:10.1111/tra.12353.

Manickam, V., A. Tiwari, J.J. Jung, R. Bhattacharya, A. Goel, D. Mukhopadhyay, and A. Choudhury. 2011. Regulation of vascular endothelial growth factor receptor 2 trafficking and angiogenesis by Golgi localized t-SNARE syntaxin 6. Blood. 117:1425–35. doi:10.1182/blood-2010-06-291690.

Mills, J., T. Hanada, Y. Hase, L. Liscum, and A.H. Chishti. 2019. LDL receptor related protein 1 requires the I3 domain of discs-large homolog 1/DLG1 for interaction with the kinesin motor protein KIF13B. Biochimica Et Biophysica Acta Bba - Mol Cell Res. 1866:118552. doi:10.1016/j.bbamcr.2019.118552.

Nakayama, M., A. Nakayama, M. van Lessen, H. Yamamoto, S. Hoffmann, H.C. Drexler, N. Itoh, T. Hirose, G. Breier, D. Vestweber, J.A. Cooper, S. Ohno, K. Kaibuchi, and R.H. Adams. 2013. Spatial regulation of VEGF receptor endocytosis in angiogenesis. Nat Cell Biol. 15:249–60. doi:10.1038/ncb2679.

Nitzsche, A., R. Pietilä, D.T. Love, C. Testini, T. Ninchoji, R.O. Smith, E. Ekvärn, J. Larsson, F.P. Roche, I. Egaña, S. Jauhiainen, P. Berger, L. Claesson-Welsh, and M. Hellström. 2021. Paladin is a phosphoinositide phosphatase regulating endosomal VEGFR2 signalling and angiogenesis. Embo Rep. 22:e50218. doi:10.15252/embr.202050218.

Orsenigo, F., C. Giampietro, A. Ferrari, M. Corada, A. Galaup, S. Sigismund, G. Ristagno, L. Maddaluno, G.Y. Koh, D. Franco, V. Kurtcuoglu, D. Poulikakos, P. Baluk, D. McDonald, M.G. Lampugnani, and E. Dejana. 2012. Phosphorylation of VE-cadherin is modulated by haemodynamic forces and contributes to the regulation of vascular permeability in vivo. Nat Commun. 3:1208. doi:10.1038/ncomms2199.

Sawamiphak, S., S. Seidel, C.L. Essmann, G.A. Wilkinson, M.E. Pitulescu, T. Acker, and A. Acker-Palmer. 2010. Ephrin-B2 regulates VEGFR2 function in developmental and tumour angiogenesis. Nature. 465:487–491. doi:10.1038/nature08995.

Simons, M., E. Gordon, and L. Claesson-Welsh. 2016. Mechanisms and regulation of endothelial VEGF receptor signalling. Nat Rev Mol Cell Bio. 17:611–25. doi:10.1038/nrm.2016.87.

Sinha, S., P.K. Vohra, R. Bhattacharya, S. Dutta, S. Sinha, and D. Mukhopadhyay. 2009. Dopamine regulates phosphorylation of VEGF receptor 2 by engaging Src-homology-2-domain-containing protein tyrosine phosphatase 2. J Cell Sci. 122:3385–3392. doi:10.1242/jcs.053124.

Smith, G.A., G.W. Fearnley, I. Abdul-Zani, S.B. Wheatcroft, D.C. Tomlinson, M.A. Harrison, and S. Ponnambalam. 2016. VEGFR2 Trafficking, Signaling and Proteolysis is Regulated by the Ubiquitin Isopeptidase USP8. Traffic. 17:53–65. doi:10.1111/tra.12341.

Smith, R., T. Ninchoji, E. Gordon, H. André, E. Dejana, D. Vestweber, A. Kvanta, and L. Claesson-Welsh. 2020. Vascular permeability in retinopathy is regulated by VEGFR2 Y949 signaling to VE-cadherin. Elife. 9:e54056. doi:10.7554/elife.54056.

Stenmark, H. 2009. Rab GTPases as coordinators of vesicle traffic. Nat Rev Mol Cell Bio. 10:513–525. doi:10.1038/nrm2728.

Tiwari, A., J.J. Jung, S.M. Inamdar, D. Nihalani, and A. Choudhury. 2013. The myosin motor Myo1c is required for VEGFR2 delivery to the cell surface and for angiogenic signaling. American journal of physiology. Heart and circulatory physiology. 304:H687–96. doi:10.1152/ajpheart.00744.2012.

Waters, S.B., J.R. Dominguez, H.-D. Cho, N.A. Sarich, A.B. Malik, and K.H. Yamada. 2021a. KIF13B-mediated VEGFR2 trafficking is essential for vascular leakage and metastasis in vivo. Life Sci Alliance. 5:e202101170. doi:10.26508/lsa.202101170.

Waters, S.B., C. Zhou, T. Nguyen, R. Zelkha, H. Lee, A. Kazlauskas, M.I. Rosenblatt, A.B. Malik, and K.H. Yamada. 2021b. VEGFR2 Trafficking by KIF13B Is a Novel Therapeutic Target for Wet Age-Related Macular Degeneration. Invest Ophth Vis Sci. 62:5. doi:10.1167/iovs.62.2.5.

Wessel, F., M. Winderlich, M. Holm, M. Frye, R. Rivera-Galdos, M. Vockel, R. Linnepe, U. Ipe, A. Stadtmann, A. Zarbock, A.F. Nottebaum, and D. Vestweber. 2014. Leukocyte extravasation and vascular permeability are each controlled in vivo by different tyrosine residues of VE-cadherin. Nat Immunol. 15:223–230. doi:10.1038/ni.2824.

Yamada, K.H., H. Kang, and A.B. Malik. 2017. Antiangiogenic Therapeutic Potential of Peptides Derived from the Molecular Motor KIF13B that Transports VEGFR2 to Plasmalemma in Endothelial Cells. The American journal of pathology. 187:214–224. doi:10.1016/j.ajpath.2016.09.010.

Yamada, K.H., Y. Nakajima, M. Geyer, K.K. Wary, M. Ushio-Fukai, Y. Komarova, and A.B. Malik. 2014. KIF13B regulates angiogenesis through Golgi to plasma membrane trafficking of VEGFR2. Journal of cell science. 127:4518–30. doi:10.1242/jcs.156109.

Zhou, H.J., Z. Xu, Z. Wang, H. Zhang, Z.W. Zhuang, M. Simons, and W. Min. 2018. SUMOylation of VEGFR2 regulates its intracellular trafficking and pathological angiogenesis. Nat Commun. 9:3303. doi:10.1038/s41467-018-05812-2.

